# Counting strands in outer membrane beta-barrels

**DOI:** 10.64898/2026.03.07.710327

**Authors:** Samuel Lim, Tejaswi Nimmagadda, Alaa Khamis, Daniel Montezano, Ryan Feehan, Matthew Copeland, Joanna S.G. Slusky

## Abstract

Beta-barrel structures are critical components of bacterial outer membranes, where they facilitate transport, cell signaling, antibiotic resistance, and structural integrity. A key feature of beta-barrels is their strand count, which influences pore diameter, binding site locations, and functional properties. However, because of breaks in strands and the presence of strands in periplasmic domains and plug domains, manual counting is inefficient and current algorithms do not accurately determine barrel strand count. To address this, we refined our previous beta-barrel structural assessment tool, PolarBearal, to improve strand number identification in large-scale datasets. To enhance the accuracy of barrel strand number labeling, our updated algorithm integrates three structural criteria, namely inter-residue vector angles, hydrogen-bonding distances, and strand connectivity. Using this algorithm, we labeled strand numbers for 571,760 predicted outer membrane beta-barrel structures obtained from the AlphaFold2 database. Our algorithm has 97% accuracy in strand number assignments, and the resulting dataset facilitates assessment of the homogeneity of strand counts for different types of outer membrane proteins. The strand labeling also provides insights on beta-barrel strand distribution and evolutionary patterns, supporting further research in protein structure prediction and design.

**Significance:** This work contributes an accurate, automated method for counting beta-barrel strands in bacterial outer membrane proteins.

*Methodological Impact:* The algorithm achieves 97% accuracy in strand counting, solving a technical problem that has hindered large-scale structural analysis. Previous manual methods were inefficient, and existing algorithms failed to handle the structural complexities of real beta-barrel proteins, including strand breaks and additional domains.

*Scale of Contribution:* By annotating over 571,000 predicted structures from the AlphaFold2 database, this work represents the largest systematic characterization of beta-barrel strand distributions to date. This labeled dataset provides a resource for the structural biology community.

*Broader Applications:* This tool enables researchers understand structure-function relationships in outer membrane proteins, supporting advances in protein design, drug development targeting bacterial membranes.

## 1. Introduction

Beta-barrel proteins are essential components in the outer membranes of mitochondria and chloroplasts, as well as in the outer membranes of Gram-negative bacteria, where they are virtually (*1*) the only structure for transmembrane proteins (*2*). In bacteria, outer membrane beta-barrels play many key roles (*2, 3*), for example: nutrient import (*4*), protein cleavage (*5*), cell signaling (*6*), toxin and antibiotic export (*7*), and cell structural integrity (*8*). Beta-barrels are all structurally similar to each other with an up-and-down meander topology. Barrels vary size by differing in the number of beta-strands, generally varying from 8 – 26, though some multi-chain barrels are larger (*9*). Because the barrel circumference is composed of a β-sheet of side-by-side strands, the strand count (also known as strand number) is generally linearly correlated with the diameter of the pore (*10*). Pore diameter is therefore a function of the number of strands, and strand count controls both the location of the loops for binding/enzyme activity and the space that can be used for passing nutrients and toxins through these proteins. Beta-barrel strand count has also been used to understand the evolutionary pathways of this fold. (*10–12*)

One of the major challenges in beta-barrel structural feature identification is accurately assigning strand numbers. Previous methods estimated this number from sequence (*13–18*), but with current higher resolution structure prediction methods (*19*) there is the possibility for improved barrel strand number estimation. However, the structures themselves do not immediately identify the barrel strand count without time-consuming visual inspection. Simple counting of a protein’s beta strands leads to counting inconsistencies in almost every structure. Because secondary structure assignment is imperfect, it can be difficult to identify the exact barrel strand number due to breaks within a strand and strands involved in extracellular or plug domains. To effectively capitalize on the new availability of predicted structures, we developed a barrel strand counting feature for our previous beta-barrel structural assessment tool, PolarBearal (*20, 21*). This updated tool identifies beta-barrel domains and strand numbers from large datasets of protein structure models. Our updated algorithm integrates multiple structural criteria to improve the accuracy of strand number assignments, generating a large-scale dataset of beta-barrel structures with barrel domain identification and high-confidence strand labels. This dataset serves as a foundation for various analyses, including evolutionary comparisons that can reveal how these structures have diversified using Evo-velocity (*22*). By refining the prediction process of counting strands and compiling a comprehensive database of beta-barrels, the updated PolarBearal program provides a valuable resource for research in better understanding the structure and function of the membrane beta-barrel on a large scale.

## 2. Methods

### 2.1 Strand identification

We used five steps to identify beta-barrel domains. Step 1: We identified potential beta-strand segments based on two structural criteria. Specifically, (a) segments must exhibit a contiguous series of beta residues with at least two consecutive residues conforming to beta structure while maintaining (b) an inter-residue angle threshold of 71 degrees or less (Figure 1A). This initial step results in the identification of individual beta-strand segments. The inter-residue angle threshold was determined in previous work to effectively differentiate between residues in barrel-forming beta strands as opposed to other common beta motifs (such as beta propellers) (*21*). Step 2: Subsequently, we merged consecutive strands that were in the same direction, specifically if the alignment of their backbone vectors was less than 80 degrees. This concatenation step combined pairs of parallel strands separated by a strand break, reducing the strand count (Figure 1C). Step 3: We enforced circular connectivity among strands, ensuring that each strand interacts with at least two other strands (Figure 1B). Step 4: Next, we eliminated residues that lack hydrogen bonds with neighboring strands or do not have nearby residues within a specified distance cutoff. Step 5: Finally, we extended any strands with neighboring residues in beta conformation. Each step was implemented as a function in the PolarBearal code. Together, these additional functions provide a framework for consistent beta-barrel strand identification and refinement.

**Figure 1:**
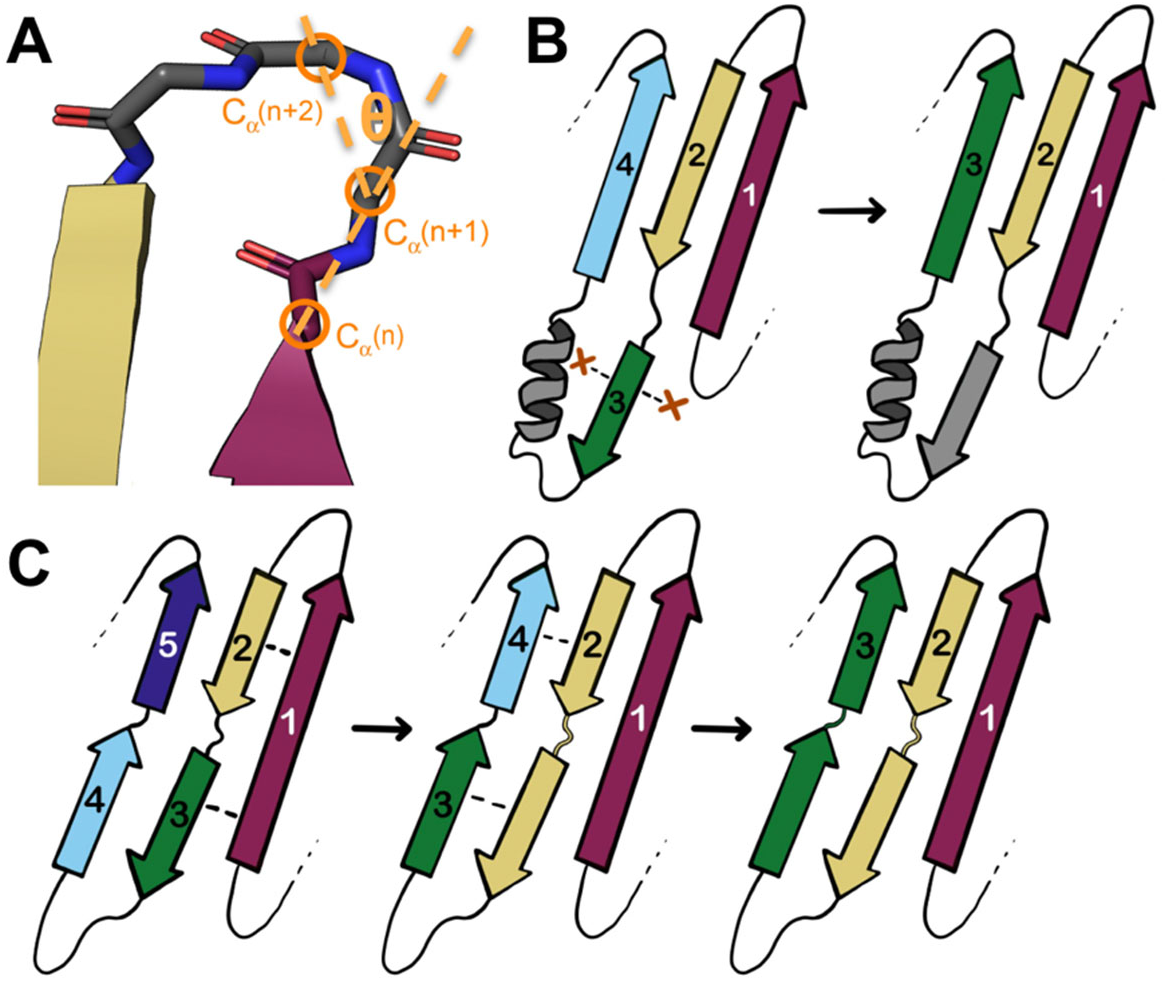
Strand counting algorithm. Strand 1 (burgundy), strand 2 (yellow), strand 3 (green), strand 4 (light blue) and strand 5 (indigo) are shown. (A) The ending of the beta-strand is determined through inter-residue vector angles formed by successive carbon alpha atoms shown in orange. (B) Removal of non-barrel beta strands is determined based on the hydrogen-bonding distance of surrounding strands. Since the non-barrel strand (green in the left image) does not have at least two neighboring strands within the hydrogen-bonding distance cutoff (as marked by dashed lines and red X), the strand is removed from the strand count (right image). (C) The merging of two consecutive strands occurs when both strands proceed in the same direction and are within hydrogen-bonding distance (dashed lines) of the same neighboring strand. This process is shown twice - merging the yellow and green strands into a single yellow strand and then merging the green and light blue strands into a single green strand.

### 2.2 Predicting Strand Count

We created a dataset of outer membrane beta-barrels by focusing on predicting the most common strand numbers. We used three steps to create a large-scale dataset of barrels with annotated strand numbers (Figure 2A). We began by mapping the 1.9 million predicted bacterial outer membrane beta-barrel protein sequences from the largest outer membrane protein dataset, IsItABarrelDB (*23*) to UniProtKB. We successfully mapped IsItABarrelDB to 812,217 UniProt IDs with pre-computed AlphaFold2 structures. We then downloaded the corresponding predicted structures from the AlphaFold2 Database (AlphaFoldDB)(*19, 24*). AlphaFold2 has previously been shown to generate outer membrane beta-barrel structures with high fidelity (*25*). Using an updated PolarBearal algorithm described in the previous section, we attempted to label the strand numbers of all 812,217 predicted structures collected from AlphaFoldDB. During the labeling process, our algorithm automatically discarded structures with fewer than five strands and categorized them as “Failed” outputs. Successful annotations were further filtered to remove low-likelihood strand number predictions (i.e. predictions with odd-numbered or uncommon strand counts). These exclusions yielded a final dataset of 571,760 proteins with the following strand counts: 8, 10, 12, 14, 16, 18, 22, 24, 26.

**Figure 2:**
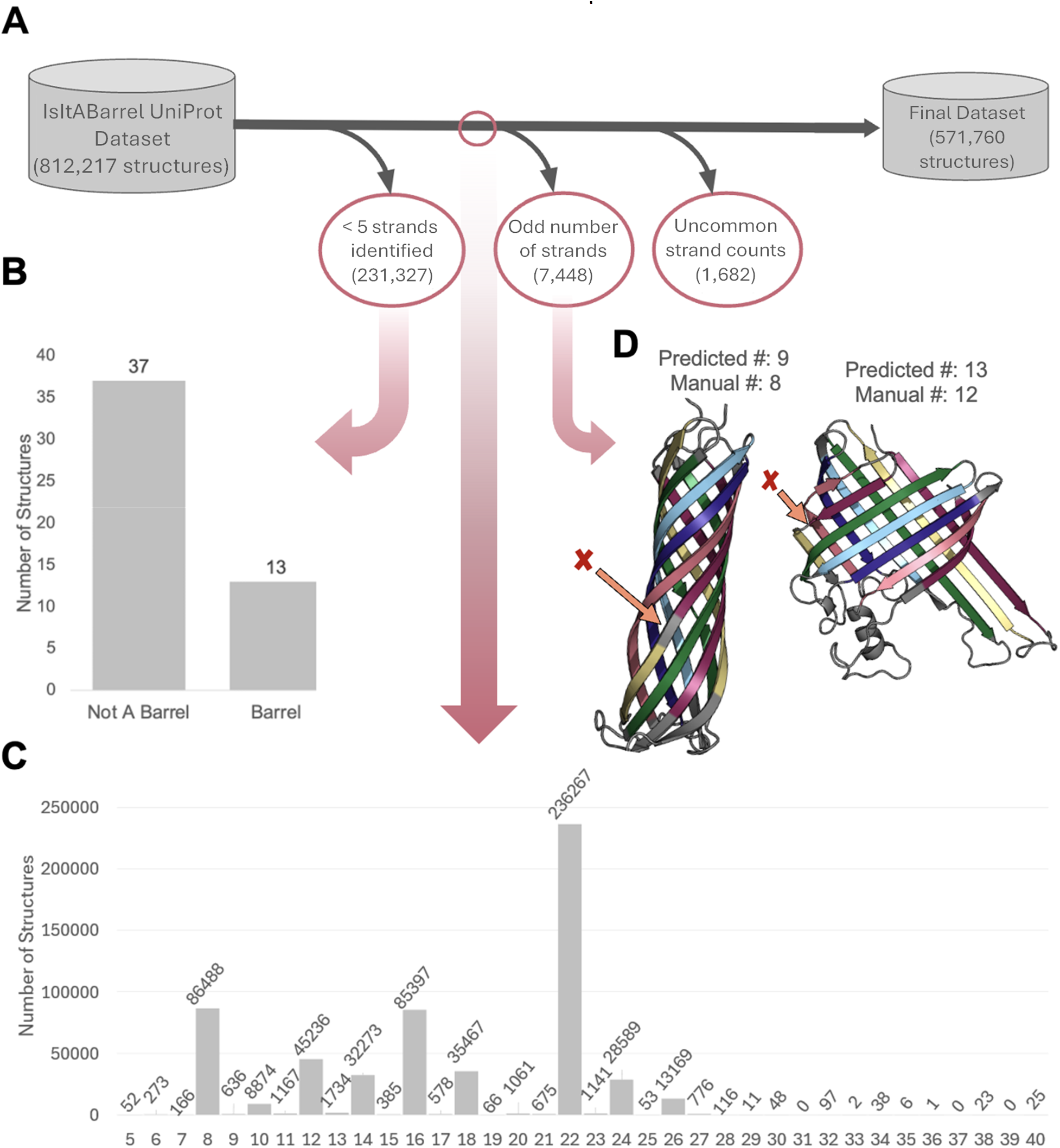
Filtering for most likely strand numbers. (A) The filtering process from the IsItABarrel UniProt dataset to the final dataset is shown through dark gray arrows pointing down to the 3 types of structures that were filtered out. The big and small gray cylinders represent the IsItABarrel UniProt dataset and the final dataset respectively. Each pink circle represents a dataset, 3 of which are analyzed in the figure as indicated by the pink arrows varying in width corresponding to the size of each dataset. (B) The manual check of 50 structures with less than 5 strands is displayed as a graph showing gray bars with 37 structures that were not barrels and 13 structures that were barrels. (C) The distribution of structures from the dataset indicated by the small pink circle in the middle of the filtering process (IsItABarrel-UniProt dataset excluding structures with less than 5 predicted strands) is shown in a barplot with the predicted strand count on the x-axis. (D) The structures from the predicted odd strand numbers dataset often resulted from unmerged strands (differentiated by color) as shown with the orange arrow with a red X indicating the location of the error. Examples are shown of 8- and 12-stranded barrels with incorrect strand number predictions.

### 2.3 Barrel Type Classification

We retrieved the barrel type classification directly from IsItABarrelDB (*23*). These barrel types are based on classifications found in the IsItABarrel database, which separated barrel sequences by similarity with sequence-similar barrels considered homologous. In that study, 11 types of sequence-dissimilar barrels were identified, and each type was found to have a different sequence motif. All barrels that did not have similarity to a type were termed “unclassified”. We used this information to explore the relationships between predicted strand count and barrel type classification in our final dataset.

### 2.4 Updating the IsItABarrel Database and PolarBearal

The predicted barrel strand numbers were updated in the downloadable and searchable IsItABarrel database and a webtool to search for specific strand numbers was implemented. This can be found at https://isitabarrel.ku.edu. The full archive is also made available for public download as a flat-file database at https://isitabarrel.ku.edu/download.

Additionally, the strand counting and barrel sizing features were updated in the downloadable and installable 3.0 version of the PolarBearal program. The algorithm and source code can be found as a Git repository at https://github.com/SluskyLab/PolarBearal3.

### 2.5 Barrel Radius as a Function of Type and Strand Number

The radius of each of the 571,760 strand-number labeled barrels were calculated as previously described using PolarBearal (*10*).

### 2.6 Evolution Inference by Barrel Strand Number

The final dataset of 571,760 beta-barrels was filtered to select only proteins from the “prototypical” barrel type (the largest of the eleven types in IsItABarrelDB), yielding a new set of 467,022 prototypical beta-barrels. This set was further reduced to a smaller set of 10,804 representative structures by eliminating redundancy with sequence clustering using MMSeqs2 (*26*). All prototypical barrels with strand labels were clustered at 30% pairwise sequence identity and 80% sequence length overlap, identical to the parameters for selecting Uniclust30 representatives, resulting in 10,804 representative prototypical beta-barrels.

We performed evolution inference analysis using Evo-velocity (*22*), a framework for inferring evolutionary dynamics from protein sequence data. Sequence embeddings were generated using the ESM1b language model. We averaged residue-level embeddings across the sequence length, to produce a per-sequence representation. The resulting embeddings were visualized using UMAP (Figure 5). Evolutionary velocities were calculated based on pseudolikelihoods to infer directional transitions in the sequence similarity network. Root and terminal points were identified as regions where velocity vectors converged or diverged using diffusion pseudotime analysis.

**Table 1.** Barrel Type Classification by Predicted Strand Number.

| Strand # | Prototypical | DUF1302 | Fim/Usher | Hypothetical Type 1 | Hypothetical Type 2 | LptD-like | LpxR-like | NfrA-like | PorT-like | TorF-like | Tsx-like | Unclassified |
| --- | --- | --- | --- | --- | --- | --- | --- | --- | --- | --- | --- | --- |
| 8 | 64316 | 0 | 0 | 0 | 501 | 0 | 0 | 0 | 1806 | 0 | 0 | 19865 |
| 10 | 3599 | 0 | 0 | 0 | 0 | 0 | 0 | 0 | 0 | 0 | 0 | 5275 |
| 12 | 19709 | 0 | 0 | 0 | 0 | 0 | 2934 | 844 | 0 | 1464 | 4557 | 15728 |
| 14 | 28598 | 0 | 0 | 0 | 0 | 0 | 0 | 5 | 0 | 0 | 0 | 3670 |
| 16 | 75742 | 0 | 1554 | 0 | 0 | 53 | 0 | 0 | 0 | 0 | 0 | 8048 |
| 18 | 30053 | 3379 | 0 | 0 | 0 | 6 | 0 | 0 | 0 | 0 | 0 | 2029 |
| 22 | 234169 | 71 | 233 | 20 | 1 | 140 | 40 | 11 | 17 | 43 | 119 | 1403 |
| 24 | 4901 | 0 | 22025 | 1436 | 0 | 4 | 0 | 0 | 0 | 0 | 3 | 220 |
| 26 | 5935 | 0 | 1 | 0 | 0 | 6789 | 0 | 0 | 0 | 0 | 0 | 444 |
| Total | 467022 | 3450 | 23813 | 1456 | 502 | 6992 | 2974 | 860 | 1823 | 1507 | 4679 | 56682 |

## 3. Results

From an initial IsItABarrel dataset, only 812,217 were mapped to UniProt IDs with precomputed AlphaFold2 structures. We predicted the number of strands for each structure (as described in Methods). We then filtered out most of the outputs which had unusual strand numbers as those had the highest levels of inaccuracy. Structures with fewer than five strands were automatically discarded, resulting in 231,327 “failed” outputs. A manual check of 50 randomly selected structures from the discarded set revealed that 37 were true negatives (not barrels), while 13 were false negatives (barrels with 5 or more strands (Figure 2B). The remaining 580,890 structures had 5 to 40 strands predicted in a barrel conformation (Figure 2C).

To better understand the barrels reported with unusual strand counts, we randomly sampled each odd strand count from 7 – 25. For all strand counts other than 11, the proteins with odd strand count labels were at least 80% miscounted even-stranded barrels or misfolded proteins by visual inspection. For example, some of the predicted odd-strand numbered structures showed split strands, where one strand was incorrectly predicted as two (Figure 2D). These miscounted strand numbers include proteins labeled as 19-stranded barrels. Although 19-stranded barrels are known to exist in mitochondria (*27, 28*), none of the bacterial proteins labeled 19-stranded had an odd strand count upon visual inspection. Notably, half of the proteins labeled 19-stranded were miscounted because they were predicted to be the previously anticipated double-barrel structure (*29*). The putative 11-stranded barrels appear to be a previously unknown fold which we will work to add to the database and characterize in a future publication. The otherwise consistent miscounting of even-stranded barrels as odd-stranded lowered our confidence that the few labeled, odd-stranded barrels are accurately predicted. Therefore, structures with odd strand count labels (7,448 structures) were filtered out. Similarly, 1,682 other proteins with low-frequency strand-count labels (6, 20, 27, 28, 29, 30, 32, 34, 35, 38, 40) were excluded, because these numbers were either due to errors in labeling or because these predictions were considered evolutionarily unlikely.

The final dataset consisted of strand counts 8, 10, 12, 14, 16, 18, 22, 24, and 26 remaining in the dataset. This collection of strand numbers represents the most likely set of barrel structures in IsItABarrelDB and serves as the foundation for subsequent analyses.

After these refinements, we find that the 571,760 structures in the final dataset are fairly high quality as measured by pLDDT, with higher structure prediction confidence for the barrel region than the rest of the protein. Only 5 structures contain a mean pLDDT below 50 (low-confidence structures, < 0.001 % of the dataset) within the barrel region (Figure S1). The overwhelming majority of our structures were found to be above a mean global pLDDT of 80 and a mean pLDDT of 90 within the barrel region, with distribution peaks at scores of 88.7 and 94.0 for both global and barrel-only mean pLDDT. (Figure S1A). 70.65% of our structures had a barrel-only mean pLDDT of 90 or higher, supporting that the data our algorithm is using is reliable. When separated by strand number, although the vast majority of barrel strand counts are highly confident in the barrel region, 8-stranded barrels have a broader distribution that extends to lower mean pLDDT scores (Figure S1B).

The final dataset of 571,760 proteins, exhibited a distribution of predicted strand counts with 22-stranded beta-barrels being the largest group (236,267 proteins), followed by 8-stranded beta-barrels (86,488 proteins) and then 16-stranded barrels (85,397 proteins) (Figure 3A). Structures with 10-stranded barrels were the least frequently predicted (8,874 proteins). To further validate the accuracy of this dataset, a manual check was performed on 100 randomly selected structures based on correct strand count and whether the models were single barrel structures. This revealed that 97 were true-positives or accurate and three were false positives or inaccurate (Figure 3B).

**Figure 3:**
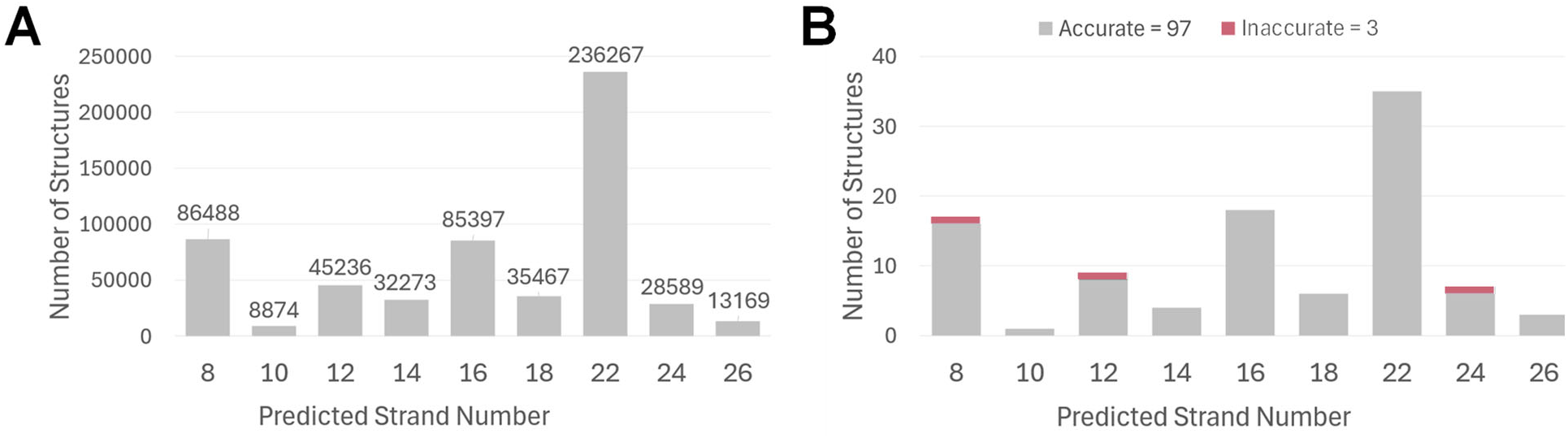
Distribution and accuracy of the Final Dataset. (A) The distribution of the final dataset is shown based on predicted strand numbers from PolarBearal. (B) The manual check on 100 structures shows inaccurate (pink) and accurate (gray) structures determined through manual strand count and whether the structure is a single barrel.

The barrel type classification of the final dataset, as documented in the IsItABarrel dataset (*23*), revealed distinct distributions across predicted strand numbers (Figure 4, Table 1). Prototypical barrels were present across all strand counts, with notable peaks at 8, 16, and 22 strands.

**Figure 4:**
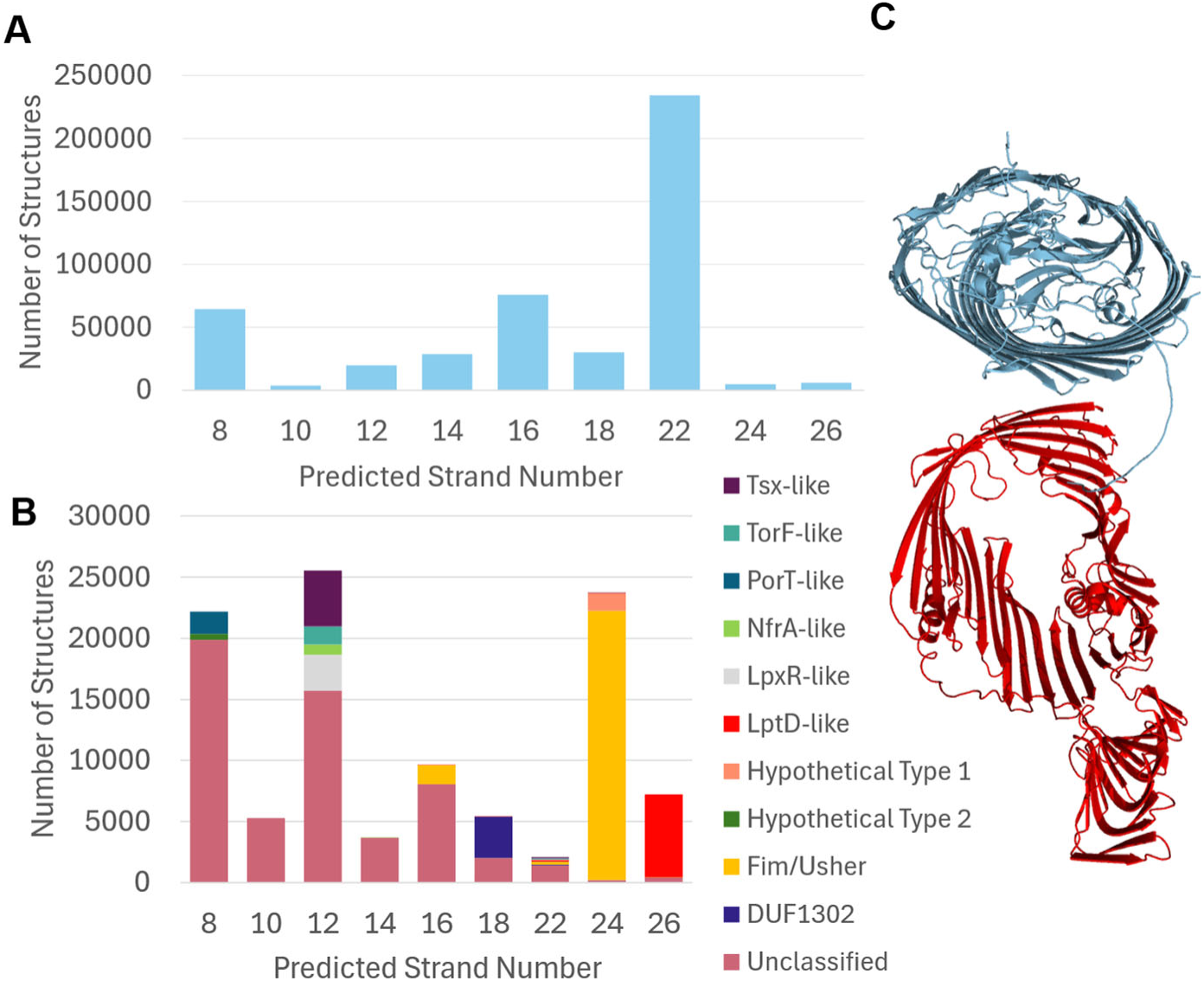
Barrel Type Classification of the Final Dataset. (A) The distribution of predicted strand counts for prototypical barrels. (B) The distribution of barrel types is shown for each predicted strand number. Details of barrel type classification shown in the key. (C) A comparison of the different geometries between the common prototypical 26-stranded barrel geometry (top, light blue) and an LptD-like barrel geometry (bottom, red).

Due to the prototypical barrels dominating the dataset (> 80%) we had to separate prototypical barrels (Figure 4A) from non-prototypical barrels (Figure 4B) so that the non-prototypical barrel features could be visualized. We find PorT-like barrels at 8 strands, Tsx-like and LpxR-like barrels at 12 strands, DUF1302 barrels at 18 strands, Fim/Usher barrels at 24 strands, and LptD-like barrels at 26 strands (Figure 4C). These findings support that specific strand counts may be indicative of functional barrel types, contributing to a broader understanding of the relationship between structure and function in beta-barrels.

Our large-scale strand count labeling can be used to better understand the differences among the barrel types. We previously found that with the exception of efflux pump proteins, outer membrane beta-barrel radius linearly correlates with strand number (*10*) We find that this remains true with this larger dataset (Figure 5A), at least for barrels with strand numbers between 8 and 18. The mean of the radii for each strand number fits well to the linear equation *y = 0.96x + 0.32 (r = 0.99).* However, for 22 - 26 stranded barrels this prediction begins to overestimate the mean radius by ≥1Å, indicating that these barrels are more oblong or kidney-shaped.

**Figure 5:**
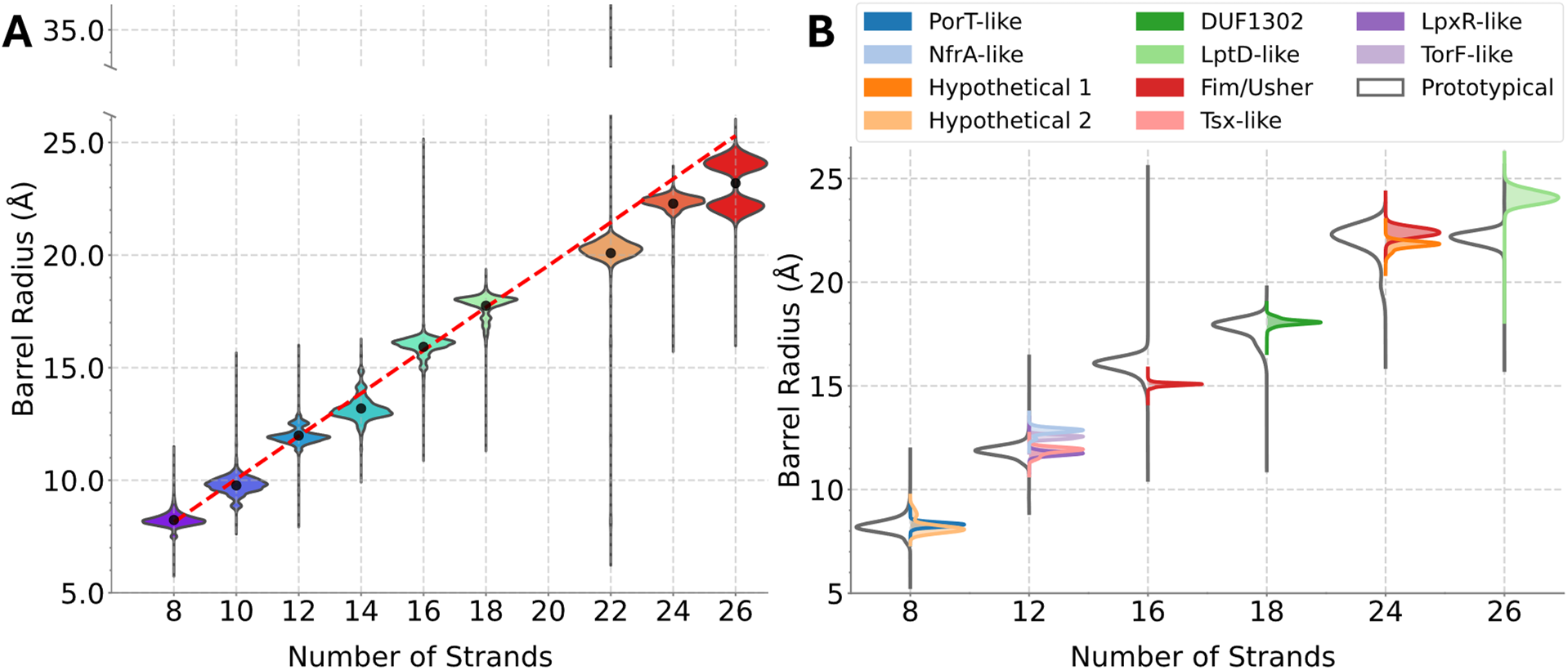
Barrel Type Distribution by Pore Size. (A) The size distribution of different strand counts by mean barrel radius. A line of best fit (dotted red) approximates the linear relationship between strand numbers 8 – 18 and barrel radius. (B) The size distribution of prototypical barrels (left, gray line) compared to each minority barrel type (colored, right) for each given strand number. Each distribution is normalized separately for illustration, though the relative quantity of each type varies (Table 1).

We also observe that each barrel type has a distinct radial distribution. Since prototypical proteins are 80% of our dataset they dominate most distributions. When we separate out the other types from the prototypical type, their radii distributions can be better compared (Figure 5B). Of the barrel classifications that make up significant minorities in each strand count ( ≥2.5% of the data for a given strand number, refer to Table 1), the majority of barrel types tend to have radii distributions within the distribution of the prototypical barrels (gray) and slightly different distribution peaks to each other. The most distinct exception is the disjoint bimodal distribution of 26-stranded barrels between LptD-like and prototypical. Tsx-like barrels (pink) and LpxR-like (purple) proteins are much closer to the size of the 12-stranded prototypical barrels while NfrA-like (baby blue) and TorF-like (lavender) predictions tend to be slightly larger. 24-stranded Fim/Usher (red) barrels have similar radial sizes to their prototypical counterparts, but 16-stranded Fim/Usher barrels are smaller than 16-stranded prototypical barrels.

Our large-scale strand count labeling algorithm can be also used to identify biological features of strand counts and relationships between strand counts, including evolutionary pathways. Recently, an evolutionary directional algorithm was developed using the protein language model ESM1b (*30*) that overcomes many limitations of traditional phylogenetic analyses (*22*). Using our dataset of beta-barrel cluster representatives with annotated strand counts we applied this algorithm to visualize strand-based evolutionary pathways (Figure 6A).

**Figure 6.**
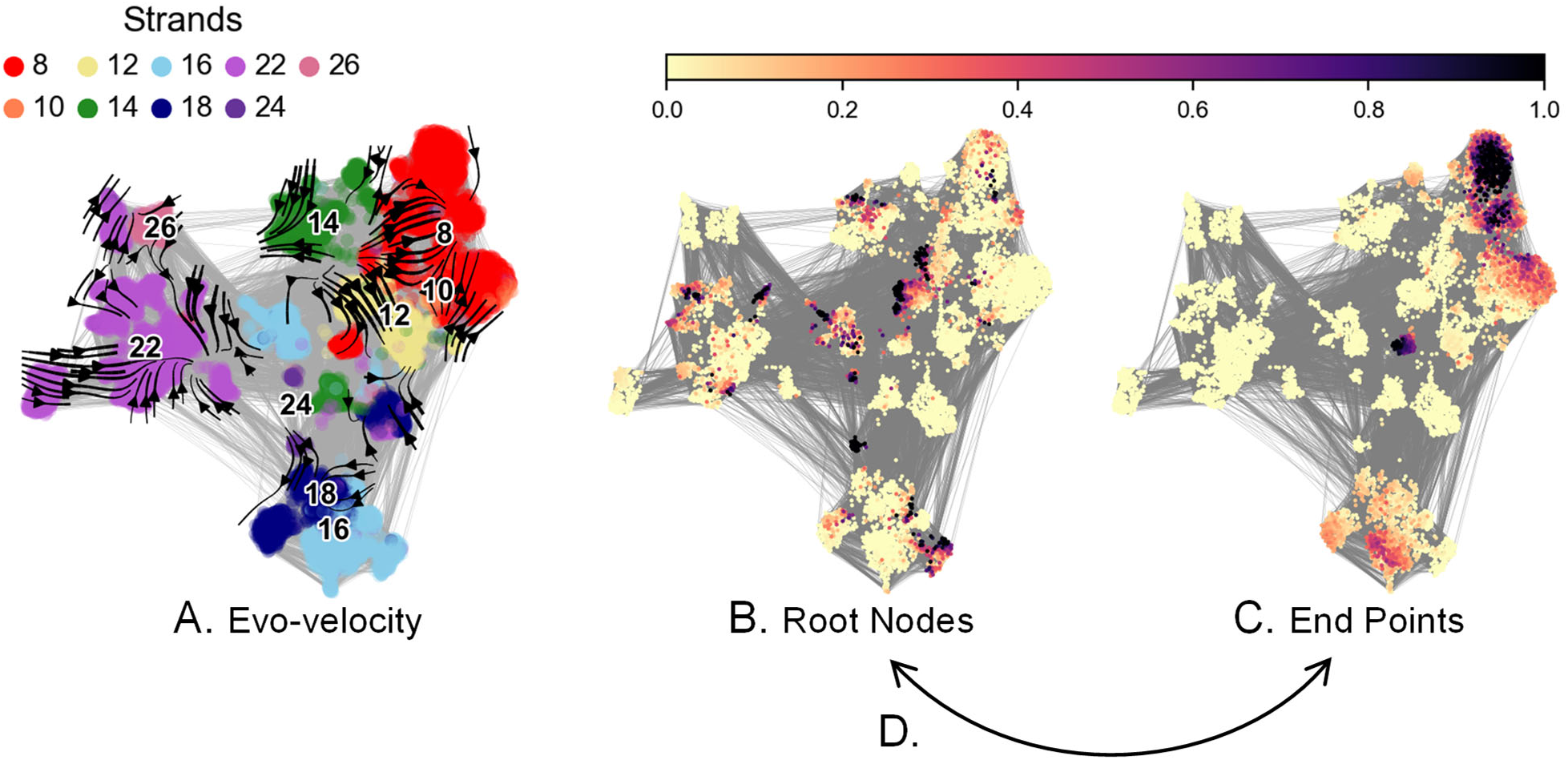
Prototypical Barrel Strand Number Evolution. For the 10,804 prototypical barrels, embeddings were visualized using UMAP with each protein represented as a node and edges showing neighbors in the high dimensional embedding space. A) with each point colored based on its protein’s strand count and computed evolutionary directional transitions in the sequence similarity network shown as arrows. B and C) Root and end nodes were calculated with pseudotime analysis. Darker colors are more likely to be (B) root nodes or (C) end points of evolution. D) Suggested swap of root node and end point labels.

In this analysis, proteins were clustered solely by sequence identity, and proteins with similar numbers of strands were not constrained to cluster together. However, similar to previous work (*11*), we find that proteins with the same number of strands tend to be more closely related, signaling a relative infrequency of strand number change. Also, 8-stranded barrels are highly related to other barrels and sit toward the center of the relational map (*11*), and 16- and 18-stranded barrels are closely related (*12*). Two notable differences are 1) the distance between 14- and 22-stranded barrels which had previously been seen to be more closely related, and 2) the presence of 26 stranded barrels which had previously not been labeled as prototypical.

Previous understanding of strand-based evolution posited a root of 8-stranded barrels diversifying to barrels with a larger number of strands (*11*). UMAP visualization of pseudotime analysis (Figure 6B and C) indicates this directionality is reversed with multiple roots at the larger barrels and end points clustered mostly at 8-stranded barrels.

## Discussion

Machine-learning-powered, accurate structure prediction has transformed protein science and created new opportunities to understand structure, function and evolution of large protein classes. However, the large influx of structures cannot instantly be translated to all forms of structural understanding. Bespoke algorithms such as strand-counting in PolarBearal can now be used to leverage these newly predicted structures. With 97% accuracy for strand number prediction and search availability via webserver, we anticipate this strand counter algorithm will support deeper insights into all beta-barrel proteins of bacterial outer membranes.

One example of a future application of our strand counting database is functional annotation. Outer membrane protein function is often tied to a barrel’s strand number— 8-stranded barrels are too small to transport solutes and so play more structural roles, 10-stranded barrels are often proteases, 16-stranded barrels are often less-specific pores, and 18-stranded barrels are often more specific pores, etc. By labeling strand numbers, we can have a better estimation of function and as functional knowledge gets better it will be more easily generalized by strand number. In this way, strand number labeling will become more useful over time.

Of the ten types of non-prototypical barrels, nine types are nearly homogenously a single strand number (Figure 5, and Table 1)— PorT-like proteins and Hypothetical Type 2 proteins are always 8-stranded, LpxR-like proteins, NfrA-like proteins, TorF-like proteins and Tsx-like proteins are 12-stranded, DUF1302 proteins are 16-stranded, hypothetical type 1 proteins are 24-stranded, and LptD-like proteins are 26-stranded barrels. Although some of these barrel types have small distributions beyond the mode, the outliers are all ≤ 2.8% of the total (TorF-like), consistent with, and exceeding, our manually assessed accuracy metric of 97%. The one truly bimodal barrel type is the Fim/Usher type which has both 16- and 24-stranded representatives. Identifying the difference in function between these two sizes of related proteins may lead to future insights of membrane structure and function diversification.

We note that although all strand numbers are represented in the prototypical barrel type, they are very much unequally distributed. Slightly over half of the prototypical barrels are 22-stranded, a strand number not used by any other barrel type. Another barrel strand number not used by other barrel types is the 10-stranded barrels, which has very little representation in the prototypicals at only 0.8%. However, this may be an under-representation of 10-stranded barrels as when double-checking the set for the most well characterized type of 10-stranded barrel, the omptin, we found that PolarBearal fails to count more than 5 strands. Specifically, the sharp vector angles between the amino acids in the strands of these structures result in failure to merge strands, and a miscounting of the strand count. This failure to accurately count omptin strands presents a limitation of the algorithm that may be overcome with future work.

While this dataset accurately represents most outer-membrane barrels, there are a few barrel topologies not represented, notably, multi-chain barrels and larger barrels. Some bacterial outer membrane barrels like TolC are assembled from symmetric protein subunits of beta-strands. Since the multimeric subunit alone does not contain a barrel, it is not found in our original dataset. Expanding the database to include these assemblies via SWISS-MODEL (*31*) or other programs may be useful to address in future work.

Another limitation of our dataset is lack of inclusion of barrels larger than 26 strands. The known proteins of these types are often either mispredicted in AFDB or too large for the automated, un-curated AFDB predictions. For example, the outer membrane protein SlpA (UniProt accession Q9RRB6) was determined by cryo-EM to be a 30-stranded barrel (*32*). However, due to its size and shallow barrel-region-MSA, the AFDB entry is misfolded and only forms a partial barrel. Such entries would be filtered out in our pipeline due to structure prediction quality. Furthermore, the largest known beta-barrel, the type 9 secretion system protein SprA (A0A2A8CX84) was determined to include a single 36-stranded barrel by cryo-EM (*33*). However, AlphaFoldDB limits un-curated predictions to 1,280 residues. Because SprA is over 2,600 residues it has no corresponding AFDB prediction and was therefore excluded from our consideration. As structure prediction technology improves, we expect that larger barrels will be more tractable to the type of analysis used here.

Although the relationships between prototypical barrel strand numbers shown here (Figure 5A) closely mirror our previous analysis with a much smaller dataset, the evolutionary hypothesis is the directional (*polar*) opposite. The evolutionary progression of strand numbers shown here supports an evolutionary start position at 22-strands as well as possible other start positions (at 8-, 10-, and 16-strands) that converge to 8-stranded barrels. This inference is in the opposite direction from our previous hypothesis which posited an 8-stranded origin diverging to other strand numbers through duplication/accretion events.

The divergence from an 8-stranded barrel hypothesis was based on two analyses. The first was a two stage strand divergence (small to mid-sized to larger) primarily through strand accretion (*11*). We calculated that 8-stranded barrels were homologous to mid-sized barrels (10-, 12-, 14-, and 16-stranded barrels) and that mid-sized barrels are homologous to even larger barrels (18- and 22- stranded) specifically due to alignments using regions of mid-sized barrels that the 8-stranded barrels align with. Although the converse direction of evolving from larger to smaller barrels was not the original hypothesis presented, it does not violate the general framework proposed.

In contrast, the second analysis is more challenging to the larger-to-smaller barrel evolutionary hypothesis. Beta-barrel proteins are repeat proteins (*34*) with a double hairpin repeat (*35*). This motif duplication in repeat proteins is thought to result from polymerase or strand slippage of DNA hairpins (*36*). Since most repeat proteins do not function as individual domains, it is believed that repeat proteins originated from ancestral homo-oligomers. Homo-oligomers would then evolve into the monomeric repeat proteins through evolutionary duplication events (*37*). Possible homomeric precursors for membrane beta-barrels include the homo-trimeric autotransporter (or their ancestors) where each subunit contributes a double hairpin (*10*). Repeat proteins are not known to start larger and then lose repeated domains, although they may delete repeated domains after initial duplication/accretion events. Therefore, it may be the case that there were initial ancient duplications from a primordial smaller barrel leading to the larger barrels—all before the modern proteins under consideration, which are subsequently diversified from larger to smaller barrels as predicted by evo-velocity. However, we view this as unlikely.

We note that due to training limitations of masked language models, the evo-velocity score is only based on substitutions, not insertions (repetitions) or deletions. This may limit evo-velocity’s utility for determining the direction of evolution of proteins that dramatically change size. Beyond evo-velocity limitations from not considering insertions and deletions, evo-velocity scoring implicitly prioritizes regions with higher sequence conservation, shorter proteins would be considered more heavily conserved by the algorithm. Therefore, evo-velocity will tend to predict the evolution of longer domains to homologous shorter ones, regardless of the true evolutionary direction. Both of these reasons support the conclusion that for repeat proteins evo-velocity’s inference of a *relational* map is more accurate than the evolutionary *directional* map. Therefore, we continue to support a model of diverging toward larger proteins over time (Figure 6D). Future work may help deconvolute which evolutionary direction is accurate and how large language models can better accommodate evolutionary predictions of repeat proteins. Large-labeled datasets, such as the one developed here, will remain valuable for these analyses.

With respect to future analyses, this dataset can provide labeled data for future analyses of outer membrane proteins, either using conventional or machine learning-based approaches. We further anticipate that this database will allow for better identification of outer membrane protein structural and functional features and how or if those features are dependent on strand number. Finally, we hope that the structural patterns identified here can be mimicked for use in designs of proteins that would be favored by nature.

## Data Availability

The full archive of barrels and those with strand counts is available for public download as a flat-file database at https://isitabarrel.ku.edu/download. Additionally, the strand counting and barrel sizing features were updated in the downloadable and installable 3.0 version of the PolarBearal program. The algorithm and source code can be found as a Git repository at https://github.com/SluskyLab/PolarBearal3. Any other data is available upon request.

## Author Contributions

SL- investigation, methodology, writing, and editing; TN- investigation, methodology, software and writing; AK- investigation, DM- methodology and editing; RF- methodology; JSGS-conceptualization, methodology, writing, and editing.

## Acknowledgements

We gratefully acknowledge Jie Liang for his contributions to the bioinformatics of outer membrane beta barrels and for his contributions as a mentor and friend to the Slusky lab. When our lab first started working on outer membrane proteins, we turned to Jie’s work the most. Jie was a gem of a colleague—always insightful, kind, and supportive. He passed too soon. We dedicate this paper to Jie’s memory. We also gratefully acknowledge support from the NSF awards MRI 2117449 and award 2226804 and the KU Self Graduate Fellowship.

## Supplementary Information

**Supplementary Figure 1:**
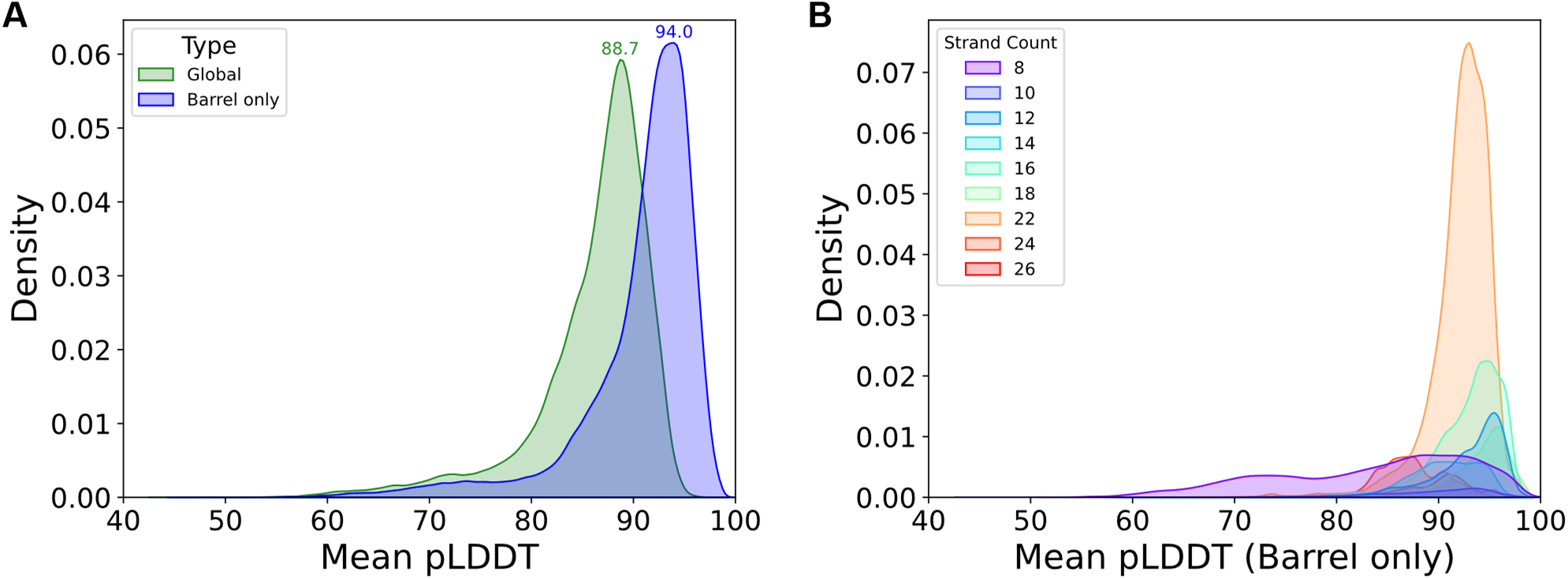
pLDDT evaluation of the final PolarBearel dataset. (A) The distributions of mean pLDDT scores measured over the whole structure (green) and for scores only in the barrel region (blue). Peak density values for global and barrel-only pLDDT confidences are 88.7 and 94.0, respectively. (B) Mean pLDDT distributions are separated by strand count. Peaks are normalized as a total percentage of the dataset, with 22-stranded barrels being the most common.

